# Machine learning reveals relationships between song, climate, and migration in coastal *Zonotrichia leucophrys*

**DOI:** 10.1101/2023.03.08.531720

**Authors:** Jiaying Yang, Bryan C. Carstens, Kaiya L. Provost

## Abstract

Vocalization behavior in birds, especially songs, strongly affects reproduction, but it is also highly impacted by geographic distance, climate, and time. For this reason, phenotypic differences in vocalizations between different bird populations are often interpreted as evidence of lineage divergence. Previous work has demonstrated that there is extensive variation in the songs of White-crowned Sparrow (*Zonotrichia leucophrys*) throughout the species range, including between neighboring (and genetically distinct) subspecies *Z. l. nuttalli* and *Z. l. pugetensis*. However, it is unknown whether the divergence in their songs correlates to environmental or geographical factors. Previous work has been hindered by time-consuming traditional methods to study bird songs that rely on the manual annotation of song spectrograms into individual syllables. Here we explore the performance of automated machine learning methods of song annotation, which can process large datasets more efficiently, paying attention to the question of subspecies differences. We utilize a recently published artificial neural network to automatically annotate hundreds of White-crowned Sparrow vocalizations across two subspecies. By analyzing differences in syllable usage and composition, we find that *Z. l. nuttalli* and *Z. l. pugetensis* have significantly different songs. Our results are consistent with the interpretation that these differences are caused by the changes in syllables in the White-crowned Sparrow repertoire. However, the large sample size enabled by the AI approach allows us to demonstrate that divergence in song is correlated with environmental difference and migratory status, but not with geographical distance. Our findings support the hypothesis that the evolution of vocalization behavior is affected by environment, in addition to population structure.

**LAY SUMMARY:**

- Birdsong is an important behavior because it is important in bird communication and reproduction.
- White-crowned Sparrows in western North America are known to use different songs along their range, but it is unknown if those songs vary due to the environment.
- We used machine learning to analyze these songs and found that populations of White-crowned Sparrows can be differentiated based on their songs.
- Environmental factors during the breeding season exert a greater influence on song evolution in migratory subspecies.

## INTRODUCTION

Vocalization is an important method of bird communication (Catchpole and Slater 2008, Brumm and Zollinger 2013), especially in *Passeriformes*. Unlike other sensory communication channels (e.g., visual, chemical, tactile), vocalizations can transmit long distances and have a fast rate of evolutionary change, due in part to cultural transmission (Slater 1986, Slater 1989, Slabbekoorn and Smith 2002a, Podos and Warren 2007, Derryberry 2009; but see Mason et al. 2017). Bird songs are vocalizations that are generally important for reproductive effort. While it was previously thought that songs were mostly produced by males during the breeding seasons, with common functions of territory defense and mating (Kroodsma and Byers 1991), more recent observations demonstrate that females also produce songs, and that songs are produced in non-breeding contexts (DeWolfe et al 1989, Baptista et al 1993, DeWolfe and Baptista 1995, Odom et al 2014, Webb et al 2016, Riebel et al 2019).

Song evolution is influenced by biotic and abiotic factors. Vocal behaviors vary among species and play a role in species recognition (Beer 1971, Emlen 1972, Baker et al. 1981b, Freeberg et al 1995). Even closely related populations can have different songs (i.e., dialects) that they can use to tell each other apart (Thompson et al. 1993). The trend of song development in birds is toward complexity and variety (i.e., the anti-monotony principle; Hartshorne 1956), mainly due to sexual selection, including male-male competition and female preference (Morton 1982, Morton 1986, Catchpole and Slater 2008). In male-male competition, listeners can determine the singer’s distance by assessing the amount of degradation of signals so that singers tend to increase the number of different song types in their repertoire (i.e., the ranging hypothesis; Morton 1982, Morton 1986).

The level of degradation and attenuation of signals is also affected by the habitat environment due to different transmitting efficacy (Acoustic Adaptation Hypothesis; Morton 1975, Hunter and Krebs 1979). Lower frequencies and slower notes transmit more effectively through denser vegetation, and birds living in those habitats have songs with those characteristics (Slabbekoorn and Smith 2002a, Naguib 2003, Kopuchian et al. 2004, Seddon 2005, Baker 2006, van Dongen and Mulder 2006, Derryberry 2009). Likewise, birds in urbanized habitats sing at higher frequencies to prevent overlap with low-frequency anthropogenic noise (Derryberry et al. 2016, Derryberry et al. 2020). Geographically restricted song variants emerge through time (Podos and Warren 2007), resulting in a pattern analogous to isolation-by-distance (Nelson et al 2004). In sum, the wide variety of bird songs likely results from any combination of sexual selection, different habitats/environments, or other biotic and abiotic factors.

### White-crowned Sparrow

One well-studied songbird is the White-crowned Sparrow (*Zonotrichia leucophrys*), which has variable song across North America (Baptista 1989). This species is known to sing year-round, not just during the breeding season, making them good candidates for understanding environmental impacts on their sounds (Blanchard 1941, Baptista 1974, Brenowitz et al. 1998). Non-breeding singing in *Z. l. pugetensis* appears to be associated with flock hierarchy formation as well as hormone stimulation (DeWolfe and Baptista 1995), in contrast with *Z. l. nuttalli* which sings due to a protracted breeding season.

White-crowned Sparrows are known to have divergent dialects across different populations (Petrinovich and Patterson 1981, Baker et al. 1984, Lien and Corbin 1990). We focus here on *Zonotrichia leucophrys nuttalli* and *Z. l. pugetensis*. These subspecies have within-species dialects, as well as divergent songs across populations associated with genetic differentiation (Lipshutz et al. 2017), suggesting that they are distinct evolutionary units. In addition, previous estimates show that dialects present in *Z. l. pugetensis* are more widely distributed than in *Z. l. nuttalli* (DeWolfe and Baptista 1995). Songs help to identify these two subspecies, both for researchers and for the birds themselves. The songs of these birds vary such that the subspecies can tell each other apart, suggesting that reproductive isolation is evolving. The subspecies have an area of hybridization in Mendocino County, CA, where intermediate individuals are found (Lipshutz et al 2017). Generally, subspecies *Z. l. nuttalli* (Figure 1A) and *Z. l. pugetensis* (Figure 1B) differ in the length of introductory whistle notes, presence of buzz notes, and length/presence of terminal trill notes (Dunn et al. 1995; Baptista and King 1980), though variation exists within populations and through time. Song is directly implicated in reducing gene flow between these populations, which have been at least partially genetically isolated for ~45,000 years (Lipshutz et al 2017). The different songs of both *Z. l. nuttalli* and *Z. l. pugetensis* also show a geographic pattern within the subspecies, which are proposed to have been formed by limited or biased dispersal, morphological variation, and/or habitat (Marler and Tamura 1962, Marler and Tamura 1964, Barker and Mewaldt 1978, Baker and Thompson 1985, Nelson 2000, Nelson et al. 2004, Kroodsma et al. 1985, reviewed by Podos and Warren 2007, Derryberry 2009).

**Figure 1.**
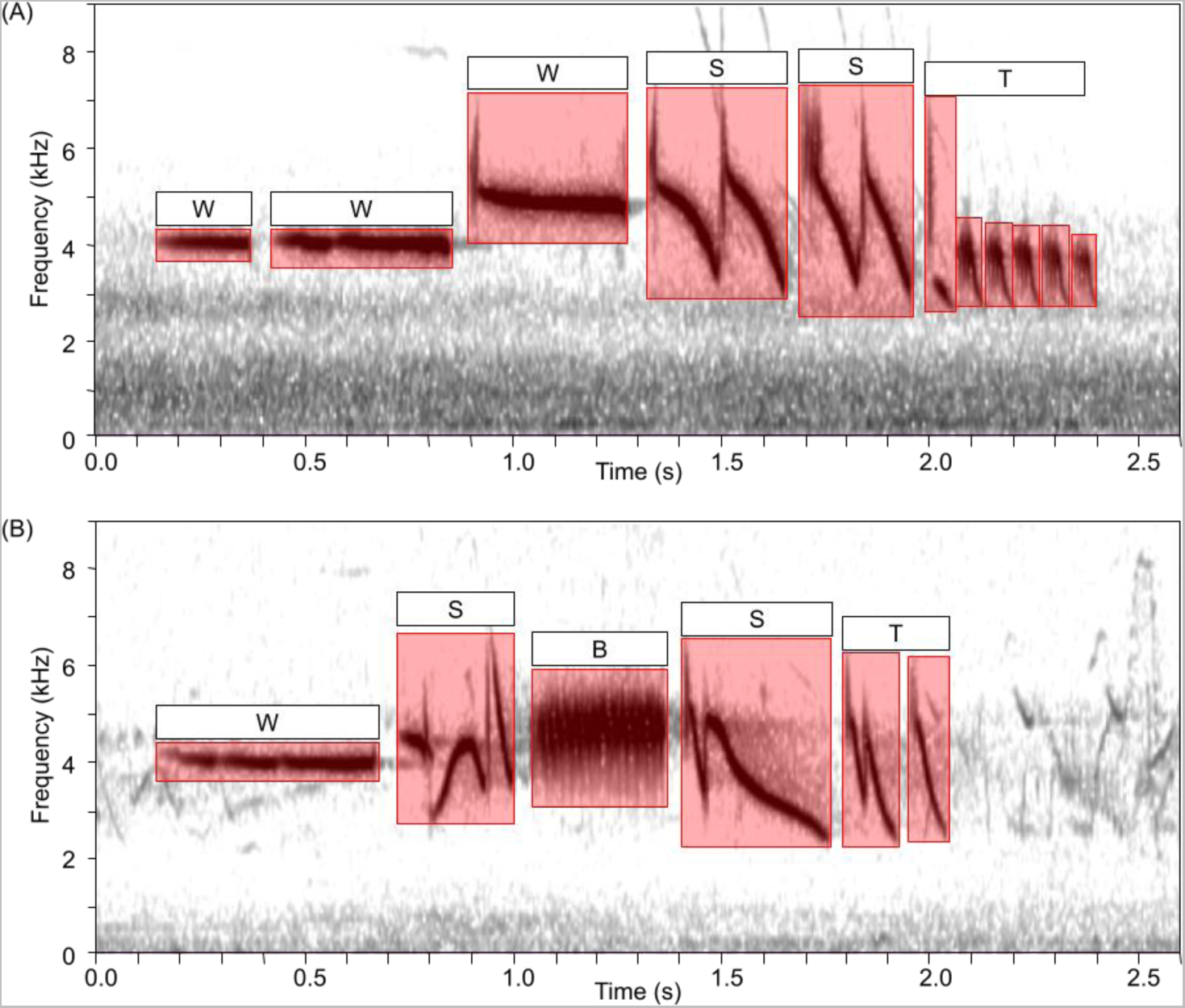
Spectrogram of (A) *Z. l. nuttalli*, BLB11552, and (B) *Z. l. pugetensis,* BLB07718, songs with manually annotation in red boxes. The X axis shows the duration time, in seconds, and the Y axis shows the frequency, in kilohertz. Higher frequencies represent higher pitch; and darker color represents higher amplitude i.e., louder sound. Each red box represents one annotated syllable, with the width representing the duration while the height indicates the differences of maximum and minimum frequency i.e., bandwidth. Note that background sounds are not annotated. Above certain notes are annotated the note type: W = whistle, B = buzz, S = “special” syllable, and T = trill of simple notes.

Though the two subspecies are phenotypically similar, with only slight differences in body size (Dunn et al. 1995), they do vary in their migratory behavior and geographic distributions. Subspecies *Z. l. nuttalli* is a non-migratory subspecies that is distributed in coastal California (from ~40.4 to 34.3 degrees latitude; Banks 1963; Figure 2). In contrast, *Z. l. pugetensis* is a migratory subspecies that is distributed from British Columbia to northwest California during the breeding season, both coastal and inland, and expands to the southern part of its range through southern California during the non-breeding season (Banks 1963, Patten et al. 2003, Baptista 1974, Heinemann 1981; Figure 2). Subspecies *Z. l. pugetensis* thus shares its wintering grounds with three other subspecies: *Z. l. nuttalli*, which is phenotypically similar, as well as *oriantha* and *gambelii*, which are distinguished by differences in color of the beak and the lores (Welke et al 2021, Lisovski et al 2019). There is evidence that White-crowned Sparrows can learn songs during migration or on the wintering grounds (in *Z. l. oriantha*, Baptista and Morton 1988). Despite this, *Z. l. pugetensis* populations having the same breeding-season dialect tend to migrate to the same location in the non-breeding season (Baptista 1974, Heinemann 1981, Nelson 2000, Nelson et al 2004) despite those locations not being segregated by song dialect (DeWolfe and Baptista 1995).

**Figure 2.**
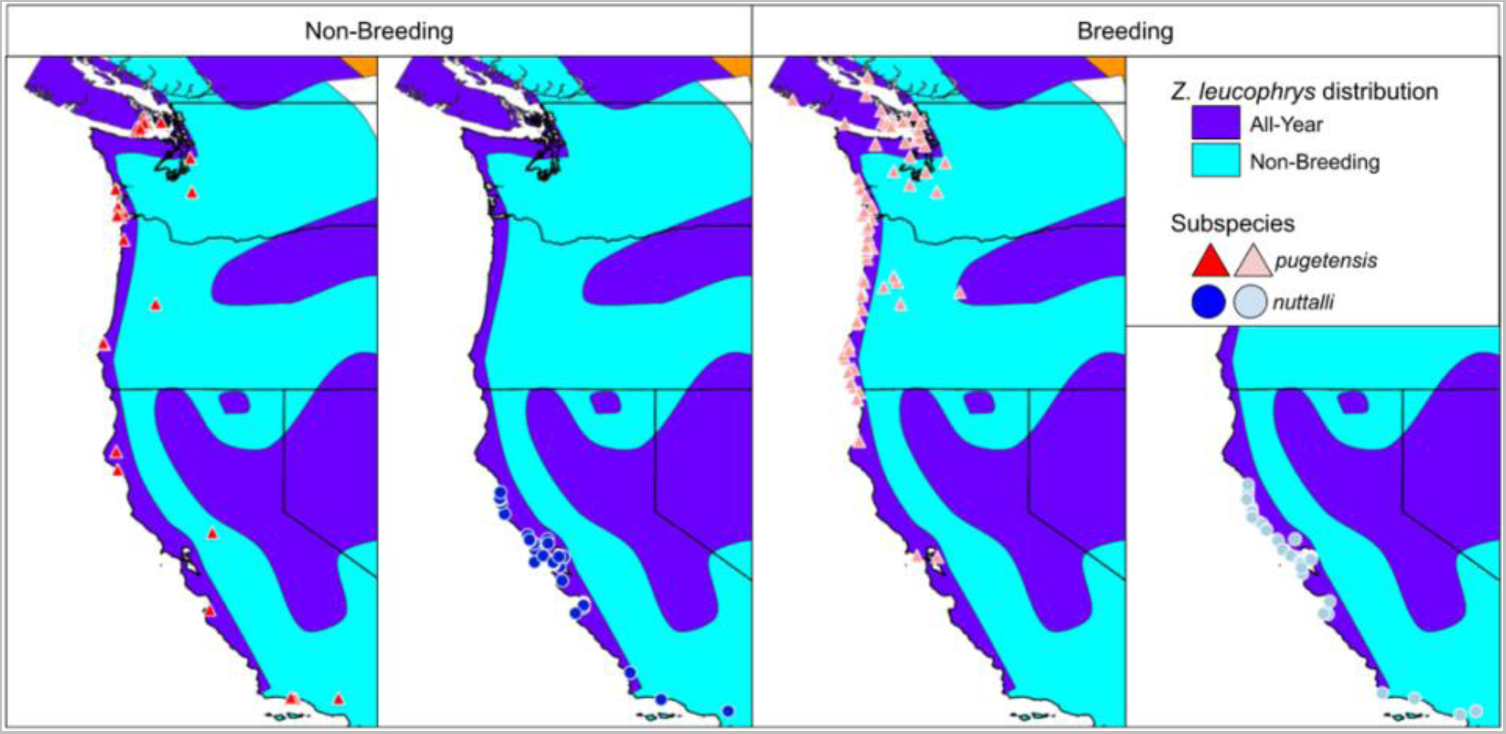
The geographical range of *Zonotrichia leucophrys* recordings that were used in this study. The sedentary *Z. l. nuttalli* individuals are blue circles and the migratory *Z. l. pugetensis* individuals are red triangles. Individuals in western California and Utah were outliers (see main text). Map below shows species distribution in all year (purple) and non-breeding season (light blue) (BirdLife International and Handbook of the Birds of the World 2021; Welke et al. 2021, Weckstein et al. 2001, Grinnell and Miller 1944, Taylor et al 2021, Cortopassi and Mewaldt 1965). Note that this map includes distributions for other subspecies (*gambelii* and *oriantha*). Individuals are separated by Non-Breeding Season (September to April, left two panels, saturated colors) and Breeding Season (May to August, right two panels, dull colors).

Though there is compelling evidence that the divergence of White-crowned Sparrow songs is caused by sexual selection (Marler and Tamula 1962, Nelson and Poesel 2013), there is less evidence on whether the evolution of vocalization behavior is correlated to the abiotic environment, such as climate, or geographic distance as a proxy for cultural evolution. Distinct differences in the histories of these two subspecies may be mediating climatic adaptation. The two subspecies overlap extensively in their non-breeding grounds. Traditionally, *Z. l. pugetensis’* breeding and wintering grounds were thought to be distinct, with the breeding not occurring south of Oregon and wintering only occurring in California (DeWolfe and Baptista 1995). However, *Z. l. pugetensis* experienced a range shift east through the Cascade Mountains and North into Alaska because of habitat change and logging regimes (Gibson et al 2013, Hunn and Beaudette 2014, Welke et al 2021). Additionally, although *Z. l. pugetensis* is assumed to migrate south to overlap with *Z. l. nuttalli*, birds identified as *Z. l. pugetensis* have been found *north* of their breeding grounds during the winter (Sullivan et al 2009; GBIF 2023). As such, it is likely that these shifts in location may be related to the adaptation of *Z. l. pugetensis* to environmental change. In this paper, we use these White-crowned Sparrow subspecies to investigate whether the divergence in songs within and between subspecies is correlated to these abiotic factors (climate and geographic distance). We hypothesize that the different songs are correlated to climates and geographical differences between the two subspecies. We also hypothesize that *Z. l. pugetensis* songs may be influenced more by the climate than that of *Z. l. nuttalli*, under the assumption that they are migratory to avoid harsh winter conditions (Somveille et al 2015), and that this impact should be more apparent during the breeding season.

### Machine learning

Bird songs have been studied for a long time with many methods for qualitatively and quantitatively describing their sounds having been developed. Traditional methods require researchers to analyze and measure song parameters manually. Since the mid 20^th^ century, the most popular methods of analysis use spectrograms, which display frequencies through time using fast Fourier transforms (Borror and Reese 1953; Cooley and Tukey 1965; Figure 1). Many analyses require every individual syllable or note on the spectrogram to be identified and annotated, or segmented, for downstream processing. Segmentation and annotation are time intensive and the amount of data that can be processed is limited (Swiston et al. 2009).

Recently, machine learning and artificial intelligence techniques have been used broadly in biological science (e.g., Schrider and Kern 2017, Pelletier and Carstens 2018, Viener et al 2022). Machine learning has also been applied to syllable segmentation with high accuracy via the neural network TweetyNet (Cohen et al. 2020). Originally trained on single species, the algorithm has since been applied to a suite of birds (Provost et al 2022) with similarly high performance, allowing for the use of larger datasets. It is unknown whether this and other automatic segmentation methods effectively recapitulate known differences in populations. Here, we examine whether we can apply automatic syllable recognition to the different songs of White-crowned Sparrow songs.

## METHODS

### Recording data and metadata

We downloaded our sound data from two sources: the Borror Laboratory of Bioacoustics (BLB; blb.osu.edu) directly from the database and the Xeno-Canto database (XC; xeno-canto.org) by using *query_xc* function in the *warbleR* package version 1.1.26 (Araya-Salas and Smith-Vidaurre 2017) in R version 4.0.3 (R Core Team 2020). We used all recordings of *Z. l. pugetensis* and *Z. l. nuttalli* in the BLB. On XC, we restricted recordings to be graded as Quality “A.” We transformed the XC recordings from MP3 format to WAV format by using the *readMP3* function in the *tuneR* package version 1.3.3 (Ligges et al. 2018) in R. We also converted recordings to have a sample rate of 48,000 Hz if needed using the *resamp* function in the *seewave* package version 2.2.0 (Sueur et al 2008) in R. Recordings were made between 1965 and 2021 and included metadata subspecies identification, date, locality, and recordist.

Although many recordings were identified to subspecies in the BLB and XC when we acquired them, both databases contained recordings from the region of interest that were not identified to subspecies. We assigned individuals without known subspecies identity using latitude and time of year as criteria following Lipshutz et al. (2017). We assigned all breeding season “unknown” individuals north of 40.544 °N and south of 48.461 °N to be *Z. l. pugetensis*, and all breeding season “unknown” individuals south of 38.419 °N and north of 37.803 °N to be *Z. l. nuttalli*, because they do not have overlapping distribution within these regions during summer. Based on the distribution, we also reassigned some mis-identified birds in the original datasets. We did not assign “unknown” individuals between the two regions because the two subspecies intergraded in that area. Moreover, “unknown” individuals that were further than ~110 km (one decimal degree) from the coast were not classified as either subspecies as they likely belonged to subspecies *Z. l. oriantha* or *Z. l. gambelii*. For non-breeding season birds, we did not assign “unknown” individuals due to substantial amounts of overlap between *Z. l. nuttalli, Z. l. pugetensis, Z. l. gambelii,* and *Z. l. oriantha* which all overwinter through the region. Any birds where subspecies identity or time of recording was not known were excluded.

Our recordings were mostly restricted to the west coast of the United States and Canada. However, there were three localities from Utah and eastern California that are in high-elevation habitats (above 1900 m), which are colder, more variable in temperature, less variable in precipitation, and drier (Fick and Hijmans 2017). As these localities were likely to influence our results, we removed them from our downstream analyses. Our final dataset included 1,111 recordings from the breeding season (547 *Z. l. pugetensis*, 564 *Z. l. nuttalli*) and 802 recordings from the non-breeding season (391 *Z. l. pugetensis*, 411 *Z. l. nuttalli*) for a total of 1,913.

### Manual Annotation

We segmented and annotated syllables present in our recordings. Segmentation is the process of going from a raw song file to a list of syllables present in the recording, including their time and frequency boundaries. Annotation is then the labeling of those syllables by syllable type (see below). For segmentation, we followed the methodology of a previous study using White-crowned Sparrows (Provost et al. 2022). 201 of these samples were segmented for that study; we manually added syllable type to those existing segmentations.

For manual segmentation and annotation, we input the recording to Raven Pro version 1.6.1 (K. Lisa Yang Center for Conservation Bioacoustics at the Cornell Lab of Ornithology 2019) and generated a spectrogram using a Fast Fourier Transform size of 512 samples. We segmented the spectrogram by drawing boxes that show the maximum and minimum values of the frequency and time of each syllable. We annotated five distinct types of syllables in these data which represent the types of syllables previously identified in *Z. leucophrys*. First, we define whistles (W) as long, single-note introductory phrases. Second, we define buzzes (B) as notes that rapidly go from high to low bandwidth. Third, we define trills (T) as individual notes that are repeated at the end of the song. Fourth, we define calls (C) as short high-pitched contact notes which are not typically used in songs. Notes that were identified as calls were excluded from downstream analyses. Lastly, we define all other notes as “special” syllables (S) if they did not fall into these categories. All annotations were output as .TXT files. We then split the original recordings and their corresponding annotation files into subsets that had approximately equal amounts of annotated syllables and silence using previously generated scripts (Provost et al. 2022).

### Using Machine Learning for Bioacoustics

To automatically segment and annotate the White-crowned Sparrow songs, we trained a TweetyNet model on our *Zonotrichia leucophrys* data following a previously described procedure (Provost et al. 2022). Although the previous study did have a model trained on *Zonotrichia leucophrys*, it was not trained on our specific syllable types from *Z. l. nuttalli* and *Z. l. pugetensis*. We used vak version 0.6.0 and TweetyNet version 0.8.0 for that and we used identical parameters to (Provost et al. 2022). We annotated 573 buzzes, 67 calls, 1,518 special syllables, 1,513 trills, and 658 whistles for 4,329 syllables. We used 80% of those as training data, 10% as validation data, and 10% as test data. After training, we discarded anything labeled as a call.

After training the model, we had the model predict the locations and annotations of syllables for all data, including data that TweetyNet had never seen before. We assessed model performance (syllable error rates, accuracy, precision, recall, and f-score) for each subspecies and per syllable type. We calculated precision and recall for each syllable type separately and then averaged them to give an overall precision and recall score.

### Summary statistics

Once we had syllables segmented and annotated for all recordings, we extracted bioacoustic data for each syllable. We extracted two types of summary statistics: one set was general to all birds, and the other was specific to White-crowned Sparrows. For the general summary statistics, data were calculated using the *spectro_analysis* function in the *warbleR* package in R. The data extracted from syllables include bandwidth (the difference between highest frequency and lowest frequency of the syllable), time duration (the length of the syllable), center frequency (central frequency between the upper and lower cutoff frequencies), average slope (slope of the change in dominant frequency), and inflection points (the local extrema). For each recording, we calculated the centroid value across all syllables. We then calculated a distance between centroids between all individuals within and between subspecies, which we used as a representation of song distance.

For the White-crowned Sparrow specific summary statistics, we calculated ten metrics that were previously identified as being important for distinguishing the two focal subspecies (Lipshutz et al 2017). These metrics are as follows: 1) the duration, 2) bandwidth, and 3) rate of the terminal trill; 4) the duration of the initial whistle notes; 5) the duration of the introductory phrase, including the initial whistle note as well as any buzzes and unique notes; 6) the maximum and 7) minimum dominant frequency present in the song; 8) the mean syllable duration, and 9) the mean dominant frequency of the first whistle note alone or 10) all initial whistle notes. We omitted metrics of vocal performance in this case.

Some of these metrics required knowledge of the position of the syllables in the recording and there were some ambiguities with how these values should be calculated. We defined the introduction as all notes before the first trill (T) note identified in the song; if not trills were identified, the introduction is the same length as the entire song. We defined the initial whistle notes as being all whistles present in the introduction if whistles were present. Finally, we defined the trill as being the duration from the first to the last trill in the song. These metrics were calculated for each song within the recording and then averaged across all songs in a single recording.

### Principal Components Analysis and ANOVAs

We performed a principal components analysis (PCA) on the vocal summary statistics present in our syllables using the *prcomp* function in the *stats* package version 4.1.2 in R. We centered and scaled all data. When values for the raw statistics were missing for the PCA specifically we filled them in with the mean value for the data. The PCA results were used to compare differences between the subspecies and seasons.

To identify the principal components (PCs) we used downstream, we used the broken-stick model (Bro and Smilde 2014) and kept all the initial PCs that explained more variation than expected under that model. We used an ANOVA on both the raw statistics and results of PCA to test the differences between the two subspecies and two seasons (breeding and non-breeding) using the *aov* function in the *stats* package in R. Given that we examined 15 raw variables (and one PC, see results) and would have a high false discovery rate across these ANOVAs, we used a Bonferroni correction (Bonferroni 1936) and set our significance value to 0.0031 from an alpha value of 0.05.

### Multiple Matrix Regression with Randomization

To determine whether differences in songs were being driven by differences in either the climate or geographic distance, we used multiple matrix regression with randomization analysis (MMRR) in R (Wang 2020). This technique allows one to interpret correlations between environmental features and geography to assess each’s relative contribution. To calculate geographic distances, we calculated the Euclidean distance between individuals with respect to latitude and longitude. For climatic differences, we downloaded 19 climate variables from WorldClim (Fick and Hijmans 2017) and extracted values at each locality in our dataset. We excluded three individuals (two *Z. l. nuttalli*, one *Z. l. pugetensis*) for whom climate information was lacking. We then calculated Euclidean distance between individuals with respect to those values. MMRR models were created such that we predicted song divergence from geographic distances and climatic differences simultaneously (i.e., song divergence ~ geographic distance + climatic distance). We also performed a second set of MMRR models where we estimate those parameters plus temporal distance, derived from the year of the recording (i.e., song divergence ~ geographic distance + climatic distance + time distance). *Zonotrichia leucophrys* is known to have experienced dramatic changes in song over the period of this study and so we wished to compare our results with and without controlling for temporal changes (Derryberry 2009). Given that we examined nine MMRR models with and without time, we used a Bonferroni correction and set our significance value to 0.0056 from an alpha value of 0.05.

Because *Z. l. pugetensis* is a subspecies that migrates through the year, we separated our data by seasons: breeding (May to August) and non-breeding (September to April; Chilton et al. 2020). We performed these MMRR analyses described above with only the breeding and non-breeding datasets separately because the different ranges due to migration of *Z. l. pugetensis* may result in variation in geographical distance and ecological distance.

### Environmental Comparisons

Given that the impact of environment on the White-crowned Sparrow songs differs between subspecies and seasons, we wished to determine whether White-crowned Sparrows in our different datasets occupy different climate regimes. We performed a PCA on the 19 WorldClim variables used in our MMRRs. We did this by extracting the climate data from all 307 unique localities that White-crowned Sparrows were found at in our dataset, and then performing a PCA on those data only. As with the PCA on song values, we used the broken stick model to select PCs to retain. We then used ANOVAs to compare whether seasons and subspecies differed in their environments. Given that we examined 19 raw environmental variables (and two PCs, see results) we used a Bonferroni correction and set our significance value to 0.0024 from an alpha value of 0.05.

## RESULTS

First, we analyzed how the machine learning model performed in syllable annotation. These models had high performance across both subspecies. Our model predicted 403 syllables on the test dataset. The syllable error rate of our trained machine learning model’s annotations was −11% for *Z. l. nuttalli* and −1% for *Z. l. pugetensis* (Table 1), such that for both subspecies, the model predicted fewer syllables than were present via manual annotation. This suggests that the models either miss some syllables outright or lump syllables together that should be separated. Although the model has a better segment error rate for *Z. l. pugetensis,* the model classifies *Z. l. nuttalli* better on all other metrics. Segmentation of the model into sounds vs background noise performed well overall (accuracy >90%, precision >96%, recall >82%, F-score>88%). Recall was the worst performing metric which was driven by the model labeling some syllables as silence. When looking at annotation of known syllables only (excluding background noise) performance is notably worse (accuracy >73%, precision >84%, recall >74%, F-score>79%). This appears to be driven largely by trills (T) which the model struggles to classify correctly (Figure 3). Trills are often misclassified as “special” (S) notes or silence.

**Figure 3:**
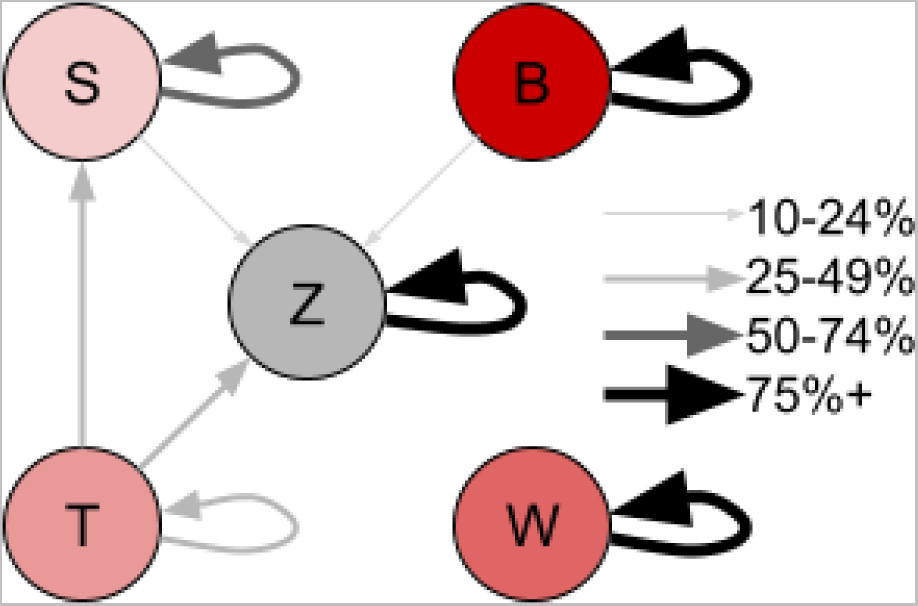
Overall syllable classifications by type. Arrows begin at the true syllable and end at the predicted syllable, with line thicknesses and color indicating the percentage of time those assignments were made. Assignments under 10% are omitted for clarity. Syllable types include buzzes (B), trills (T), whistles (W), “special” syllables (S), and zero syllables, i.e., background noise (Z)

**Table 1.** The machine learning model showed a high accuracy in annotating the syllables of both *Z. l. pugetensis* and *Z. l. nuttalli* for all data.

|  |  | Subspecies |  |  |
| --- | --- | --- | --- | --- |
|  |  | Both | *Z. l. nuttalli* | *Z. l. pugetensis* |
|  | Segment Error Rate | -6.93% | -11.43% | -1.06% |
|  | Segment Error Rate (Abs) | 6.93% | 11.43% | 1.06% |
| Sound/Background | Accuracy | 91.78% | 92.95% | 90.67% |
|  | Precision | 97.16% | 98.10% | 96.17% |
|  | Recall | 84.61% | 86.84% | 82.35% |
|  | F-Score | 90.45% | 92.12% | 88.72% |
| Syllable Type | Accuracy | 87.43% | 87.97% | 86.92% |
|  | Precision | 86.20% | 87.44% | 84.71% |
|  | Recall | 76.34% | 78.93% | 74.65% |
|  | F-Score | 80.97% | 82.97% | 79.36% |
| Syllable Type (Silence Excluded) | Accuracy | 75.15% | 76.36% | 73.91% |
|  | Precision | 85.71% | 87.00% | 84.06% |
|  | Recall | 70.96% | 74.40% | 68.97% |
|  | F-Score | 77.64% | 80.00% | 75.78% |

### PCA

We found that our PCA on syllable variables explained a high proportion of the vocal data, with the first three PCs explaining 33%, 15%, and 11% of variation, respectively (Table 2). Under the broken stick model these numbers were expected to be 22, 15, and 12%, respectively. As such, we retained only PC1 for downstream analyses. Syllables with high PC1 had low whistle dominant frequencies, few inflection points, low maximum and center frequencies, and short syllable introduction durations. They also had wide bandwidths (Table 2).

**Table 2:**
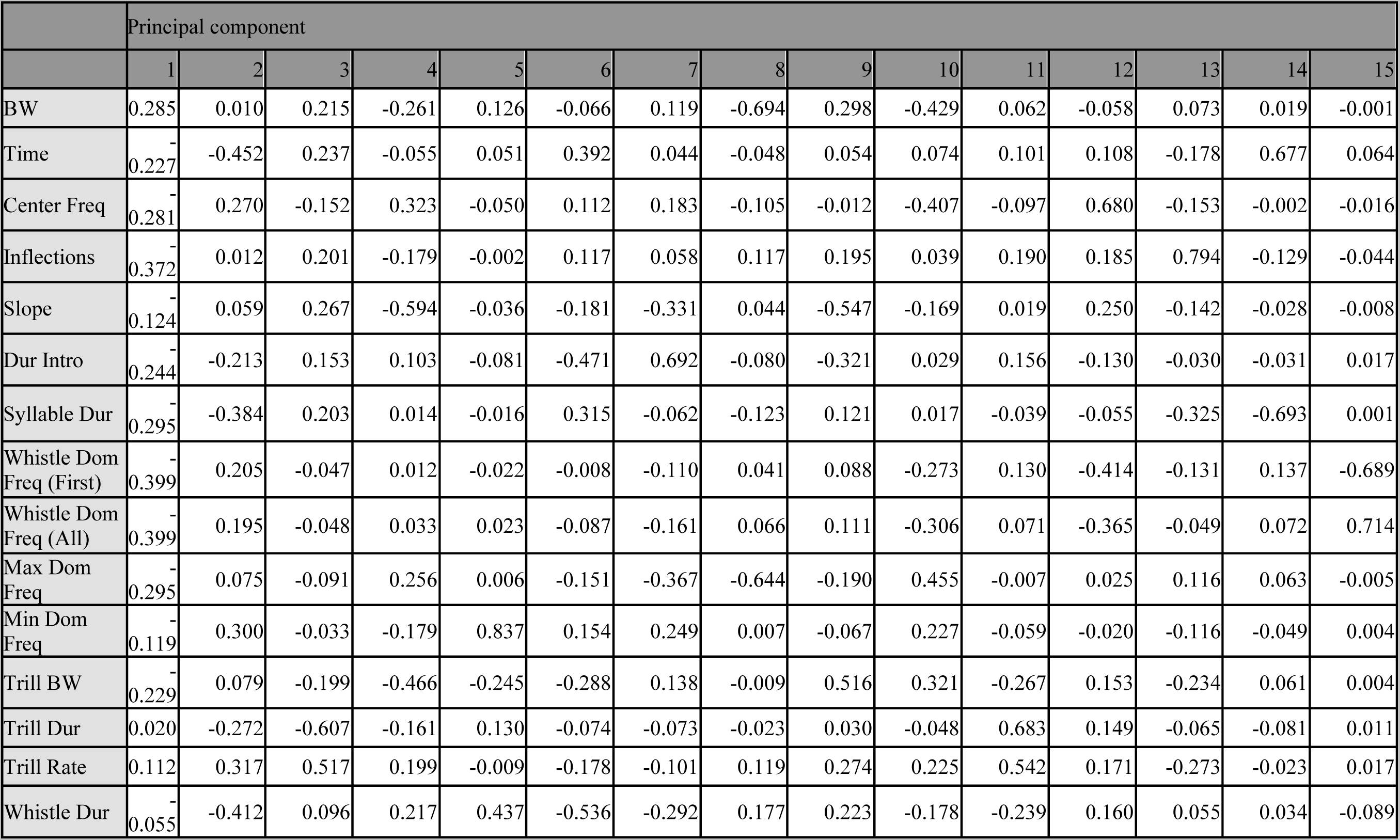
Principal component rotation of individual PCs axes for song across summary statistics of song properties. “BW” = bandwidth. “Dur” = Duration. “Dom” = Dominant. “Freq”=Frequency.

We found that our summary statistics of song did differentiate subspecies and seasons (Table 4). PC1 was significantly higher in *Z. l. nuttalli* than *Z. l. pugetensis* (p<0.0001). Most of the underlying song statistics that PC1 was calculated from also differed significantly between subspecies. Subspecies *Z l. pugetensi*s had songs with steeper slopes, wider trill bandwidths, shorter whistles, longer introductory phrases and overall syllables, higher whistle and overall song frequencies, more inflection points, and shorter and faster trills. Likewise, PC1 was higher in the breeding season than the non-breeding season, although this relationship was only nearly significant after correcting for multiple tests (p=0.0104). Despite PC1 not being significant, some of the individual statistics did show differences. Breeding season birds have narrower trill bandwidths, longer trills, lower whistle dominant frequencies but higher minimum dominant frequencies, steeper slopes, and fewer inflection points than non-breeding season birds.

**Table 3:** Principal component importance and eigenvalues of individual PCs axes for song across summary statistics of song properties. Broken stick refers to expected proportion of variance explained under the broken stick model.

| PC | Standard deviation | Proportion of Variance | Cumulative Proportion | Broken Stick |
| --- | --- | --- | --- | --- |
| 1 | 2.226 | 0.330 | 0.330 | 0.221 |
| 2 | 1.488 | 0.148 | 0.478 | 0.155 |
| 3 | 1.275 | 0.108 | 0.586 | 0.121 |
| 4 | 1.232 | 0.101 | 0.687 | 0.099 |
| 5 | 0.919 | 0.056 | 0.744 | 0.082 |
| 6 | 0.887 | 0.052 | 0.796 | 0.069 |
| 7 | 0.801 | 0.043 | 0.839 | 0.058 |
| 8 | 0.749 | 0.037 | 0.876 | 0.048 |
| 9 | 0.663 | 0.029 | 0.906 | 0.040 |
| 10 | 0.648 | 0.028 | 0.934 | 0.033 |
| 11 | 0.592 | 0.023 | 0.957 | 0.026 |
| 12 | 0.538 | 0.019 | 0.976 | 0.020 |
| 13 | 0.466 | 0.015 | 0.991 | 0.014 |
| 14 | 0.341 | 0.008 | 0.999 | 0.009 |
| 15 | 0.145 | 0.001 | 1.000 | 0.004 |

**Table 4:** Differences between season (breeding vs non-breeding) and subspecies (*Z. l. pugetensis* vs *Z. l. nuttalli*) using ANOVAs while controlling for both. Degrees of freedom = 1 for all comparisons. Note that unknown season individuals were removed. SSQ = sum of squares. MSQ = mean sum of squares. F = F-statistic. P = p-value. For these analyses, significant values have p<0.0031. We indicate a near-significant p-value with bold text and a significant value with bold text plus an asterisk.

|  | Mean values |  |  |  | Model parameters: season |  |  |  | Model parameters: subspecies |  |  |  |
| --- | --- | --- | --- | --- | --- | --- | --- | --- | --- | --- | --- | --- |
|  | breeding | non-breeding | nuttalli | pugetensis | SSQ | MSQ | F | P | SSQ | MSQ | F | P |
| PC1 | 0.255 | 0.01 | 0.874 | -0.59 | 29 | 29.3 | 6.57 | **0.01** | 1037 | 1036.7 | 232.55 | **0\*** |
| BW | 2360 | 2390 | 2380 | 2360 | 585200 | 585171 | 0.679 | 0.41 | 320900 | 320875 | 0.372 | 0.542 |
| Center Freq | 3510 | 3440 | 3330 | 3630 | 2036000 | 2036430 | 3.083 | 0.0793 | 41240000 | 41238170 | 62.425 | **0\*** |
| Inflections | 8.63 | 9.36 | 7.36 | 10.5 | 268 | 268 | 11.1 | **0\*** | 4672 | 4672 | 193.3 | **0\*** |
| Dur Intro | 1.28 | 1.27 | 1.16 | 1.41 | 0.01 | 0.008 | 0.05 | 0.824 | 22.91 | 22.909 | 135.53 | **0\*** |
| Syllable Dur | 0.187 | 0.189 | 0.18 | 0.197 | 0.002 | 0.0022 | 0.606 | 0.436 | 0.138 | 0.13778 | 37.884 | **0\*** |
| Slope | -40.7 | -30.3 | -47.9 | -24.5 | 51135 | 51135 | 23.95 | **0\*** | 253743 | 253743 | 118.85 | **0\*** |
| Whistle Dom Freq (First) | 1.86 | 2.11 | 1.49 | 2.49 | 31.1 | 31.1 | 24.58 | **0\*** | 444.4 | 444.4 | 351.43 | **0\*** |
| Whistle Dom Freq (All) | 2.06 | 2.28 | 1.73 | 2.61 | 25 | 25 | 21.31 | **0\*** | 343.6 | 343.6 | 293.16 | **0\*** |
| Max Dom Freq | 5.21 | 5.33 | 4.93 | 5.63 | 7 | 7.22 | 2.458 | 0.117 | 219 | 218.64 | 74.428 | **0\*** |
| Min Dom Freq | 0.184 | 0.138 | 0.169 | 0.159 | 0.9 | 0.9143 | 5.039 | **0.024** | 0 | 0.0409 | 0.225 | 0.6351 |
| Time | 0.21 | 0.21 | 0.208 | 0.212 | 0 | 0.000015 | 0.003 | 0.958 | 0.01 | 0.009564 | 1.801 | 0.18 |
| Trill BW | 0.0017 | 0.002 | 0.0015 | 0.0022 | 0.00003 | 0.00003 | 27.27 | **0\*** | 0.0001676 | 0.0001676 | 147.47 | **0\*** |
| Trill Dur | 0.244 | 0.228 | 0.264 | 0.206 | 0.11 | 0.1106 | 4.617 | **0.031** | 1.3 | 1.3015 | 54.319 | **0\*** |
| Trill Rate | 10.2 | 10.2 | 10 | 10.5 | 0 | 0.11 | 0.011 | 0.915 | 76 | 75.99 | 7.642 | **0.006** |
| Whistle Dur | 0.562 | 0.552 | 0.602 | 0.51 | 0.06 | 0.06 | 1.559 | 0.212 | 3.8 | 3.803 | 99.624 | **0\*** |

### MMRR

We found that geography explained song divergence for both species together, irrespective of season (R² > 0.01, geography p<0.001; Table 5; Table 6). Climate is a significant factor during the breeding season and year-round (p<0.001), but only nearly significant during the non-breeding season (p=0.013). After we account for temporal changes, these patterns are identical, but year of recording is also significant (R² > 0.05, time p<0.001, climate p<0.016, geography p<0.001).

**Table 5:**
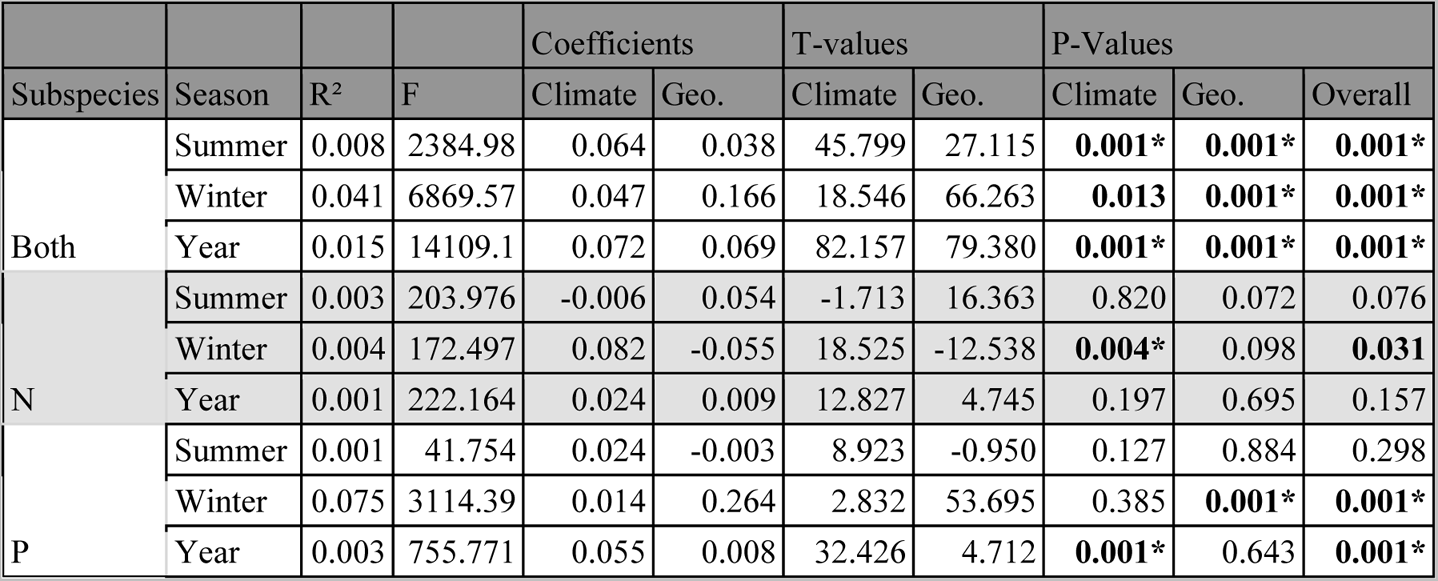
MMRR analysis shows that divergence in songs of *Z. l. pugetensis* and *Z. l. nuttalli* is strongly correlated with environmental differences before accounting for time. For these analyses, significant values have p<0.0056. We indicate a near-significant p-value with bold text and a significant value with bold text plus an asterisk.

**Table 6:** MMRR analysis shows that divergence in songs of *Z. l. pugetensis* and *Z. l. nuttalli* is strongly correlated with environmental differences even after accounting for time. For these analyses, significant values have p<0.0056. We indicate a near-significant p-value with bold text and a significant value with bold text plus an asterisk.

| Subspecies | Season | R<sup>2</sup> | F | Coefficients |  |  | T-values |  |  | P-Values |  |  |  |
| --- | --- | --- | --- | --- | --- | --- | --- | --- | --- | --- | --- | --- | --- |
|  |  |  |  | Time | Climate | Geo. | Time | Climate | Geo. | Time | Climate | Geo. | Overall |
| Both | Breeding | 0.114 | 26395 | 0.327 | 0.039 | 0.031 | 271.742 | 29.514 | 23.202 | **0.001\*** | **0.001\*** | **0.001\*** | **0.001\*** |
|  | Non-breeding | 0.052 | 5817 | 0.110 | 0.042 | 0.129 | 59.679 | 16.783 | 50.211 | **0.001\*** | **0.016** | **0.001\*** | **0.001\*** |
|  | Year | 0.073 | 47858 | 0.244 | 0.048 | 0.046 | 337.048 | 56.164 | 54.598 | **0.001\*** | **0.001\*** | **0.001\*** | **0.001\*** |
| N | Breeding | 0.050 | 2764.570 | 0.223 | -0.070 | 0.077 | 88.688 | -21.033 | 23.624 | **0.001\*** | **0.009** | **0.007** | **0.001\*** |
|  | Non-breeding | 0.010 | 269.207 | 0.080 | 0.048 | -0.047 | 21.464 | 10.278 | -10.738 | **0.001\*** | 0.092 | 0.148 | **0.007** |
|  | Year | 0.022 | 3463.662 | 0.149 | -0.024 | 0.027 | 99.686 | -12.232 | 14.219 | **0.001\*** | 0.217 | 0.231 | **0.001\*** |
| P | Breeding | 0.133 | 7589.539 | 0.364 | 0.013 | -0.010 | 150.574 | 5.101 | -3.790 | **0.001\*** | 0.364 | 0.575 | **0.001\*** |
|  | Non-breeding | 0.076 | 2097.494 | 0.037 | 0.012 | 0.240 | 7.680 | 2.473 | 40.915 | 0.342 | 0.447 | **0.001\*** | **0.001\*** |
|  | Year | 0.088 | 14167.874 | 0.293 | 0.025 | -0.002 | 202.118 | 15.334 | -1.236 | **0.001\*** | **0.010** | 0.899 | **0.001\*** |

For *Z. l. nuttalli* alone, we found a relationship between climate and song in the non-breeding season (R² = 0.004, climate p=0.004, geography p=0.098). However, this relationship is not present when accounting for time, suggesting that year of recording explains this pattern (R² = 0.010, time p=0.001, climate p=0.092, geography p=0.148). There was no impact of geography or climate on song for *Z. l. nuttalli* during the breeding season. After accounting for time, there is an association during the breeding season between song and time, but climate and geography are both nearly significant (R² = 0.05, time p=0.001, climate p=0.002, geography p=0.007). In contrast for the full year, while there was no impact of geography and climate on song before accounting for time, time itself does significantly explain song.

As for *Z. l. pugetensis*, in the non-breeding season there is an association between song and geography (R² = 0.075, climate p=0.385, geography p=0.001); this association holds up after accounting for time, but time is not a significant predictor. Year-round there is an association between song and climate (R² = 0.001, climate p=0.001, geography p=0.643), but this association is only near significant after accounting for time, with time instead explaining the variation. Finally, while there was no impact on *Z. l. pugetensis* song in the breeding season before accounting for time, after accounting for time both time and climate are significant. In all instances where year of recording significantly explains song divergence, it is likely that this is reflecting patterns of cultural evolution caused by young sparrows learning their songs from adults near them (Baptista and Morton 1988).

### Environmental Comparisons

The first three environmental PCs (Env.PCs) explained 63%, 21%, and 10% of the environmental variation, respectively (Supplementary Material Table S1). Under the broken stick model, these should have explained 18%, 13%, and 11%, respectively. As such we retained the first two PCs. High values of Env.PC1 indicated environments that were cold, wet, and had little seasonal precipitation variation (Supplementary Material Table S2, Supplementary Material Table S3). High values of Env.PC2 were associated with temperature, specifically environments that had elevated temperature seasonality and annual range, and little isothermality.

ANOVAs revealed that environments were significantly different between seasons and subspecies (Supplementary Material Table S4, Supplementary Material Table S5). Subspecies significantly differ across the Env.PC1 and Env.PC2, with *pugetensis* occupying colder, wetter, and more variable areas. Seasons also differ in Env.PC1 such that breeding season birds are on average in areas that are more seasonal and less isothermal. These patterns likely reflect differences in where individual birds are located during the breeding and non-breeding seasons, especially as *pugetensis* migrates south.

## DISCUSSION

Our analysis demonstrates that the trained machine learning model by TweetyNet has high accuracy with low error rates and can annotate *Z. l. nuttalli* and *Z. l. pugetensis* songs. Our results suggest that the annotation by the machine learning model is as reliable as manual annotations and produces comparable data, although training with more diverse syllable types may improve the classification of our models, particularly on trill syllables. Given its computational efficiency, machine learning can be a useful tool to automatically annotate bird songs and extract data because it can be used to process large datasets quickly and accurately.

We found significant differences in the syllables used in the songs of *Z. l. nuttalli* and the songs of *Z. l. pugetensis*, which corroborates previous evidence that these two taxa have divergent songs (e.g., Lipshutz et al. 2017). Our bioacoustic metrics found that on average, *Z. l. nuttalli* syllables were lower in pitch, had fewer inflections, and had more negative slopes compared to *Z. l. pugetensis,* as well as differing in trill duration and bandwidth. Our analyses cannot distinguish if this is due to subspecies using different syllables or the same syllables at different frequencies. Preliminary analyses of syllable shape suggest that the differences are mediated using different syllables, but further investigation is needed to confirm.

We also found significant differences between the seasons in some metrics. Syllables used in the breeding season differed in trill bandwidth, whistle dominant frequency, slope, and inflections compared to the non-breeding season. In other species of songbird, singing in the non-breeding season is less consistent due to differences in hormone levels (e.g., the Song Sparrow *Melospiza melodia*, Smith et al 1997; the Rufous-collared Sparrow *Zonotrichia capensis*, Addis et al 2010), which is consistent with our findings. At the same time, this species modulates its song with respect to the environment, being especially impacted by noise transmission (Derryberry 2009, Luther et al 2016, Derryberry et al 2020). Additionally, distinct parts of *Zonotrichia leucophrys* song appear to have distinct functions (Nelson and Poesel 2007): introductory whistles (as well as buzzes) are highly conserved and serve as species identifiers, terminal trills are moderately conserved and indicate dialect, and other notes in the song are highly variable and indicate individual identity. Whether this species is responding to their environments in different seasons or simply singing less consistently due to a lack of testosterone, there is clearly detectable variation associated between the seasons and the environment.

The differences in song between subspecies and seasons are compounded by the fact that the two subspecies have different responses to climate, geographic distance, and time. Almost every analysis showed that year of recording partially or fully predicted variation in song (except for non-breeding season *Z. l. pugetensis*). Thus, the impact of cultural drift on this taxon cannot be overstated and corroborates previous analyses (Derryberry 2009). Geographic distance is important when both subspecies are analyzed simultaneously, regardless of season. Additionally, the migratory *Z. l. pugetensis* also shows a high impact of geographic distance on songs recorded in the non-breeding season. In contrast, there is mixed evidence whether geographic distance explains song variation in the sedentary *Z. l. nuttalli.* The impact of climate on song is important when both species are analyzed together year-round and during the breeding season. In contrast, climate does not explain variation in either subspecies after accounting for time. The evidence suggests that while cultural drift through time is important for these two taxa in explaining song differences, overall, the migratory subspecies *Z. l. pugetensis* shows higher influence of spatial factors, particularly geographic distance, though only during the non-breeding months. This may be driving the patterns where both species are combined in those seasons; alternatively, the two subspecies may still be adapted to their relatively different environments. It is likely that these factors (i.e., migration and seasonality) have contributed to the evolution of song differences.

We performed an analysis of the environments that each of the subspecies occupy during different seasons and found that these climates are significantly different. Year-round, *Z. l. nuttalli* occupies warmer and drier areas than *Z. l. pugetensis* does in the summer. In contrast, wintering *Z. l. pugetensis* experience nearly the full yearly range of temperature and precipitation across both subspecies, and overall experience more variable and more seasonal climates than *Z. l. nuttalli*. A synthesis across multiple birds hypothesizes that migration occurs as a mechanism to avoid harsh winters and exploit resource surpluses (Somerville et al 2015); given that the range of *Z. l. pugetensis* is wide-spread and has been changing in response to habitat disruption (Welke et al 2021, Hunn and Beaudette 2014, Gibson et al 2013), we hypothesize that *Z. l. pugetensis* as a subspecies may be more susceptible to spatial heterogeneity in their environment, leading to more variation in their songs. In addition, because *Z. l. pugetensis* is known to have dialects ranging over larger areas that encapsulate more variable climates (DeWolfe and Baptista 1995), it is possible that the selection pressures to adapt song to environment is higher across the range of this subspecies.

### Caveats

Several caveats are worth documenting to place our results in the appropriate context. All analyses suggest that geographic distance is not correlated to the divergence in song across these two subspecies, which is not consistent with our hypothesis. Some explanations are as follows. First, although the number of recordings of *Z. l. pugetensis* in non-breeding habitats is large, they do not cover the entire projected breeding range. Second, our recordings are all from the BLB and XC database from 1965 to 2021, resulting in substantial cultural drift. Over the time scale of the study, White-crowned Sparrows are known to have demonstrably changed their songs (e.g., Derryberry 2009). However, even if we control for the year of recording, we still find significant relationships with spatial factors like geography and climate. Third, our analysis only used each syllable’s characteristics and types but did not analyze the syntax or order of syllables. It is possible that they used the same syllables but different orders in some populations. Fourth, because White-crowned Sparrows learn their songs not only directly from their parents but also from the neighboring community (Cunningham and Baker 1983) and that individuals in hybrid zones can learn the songs of the “incorrect” subspecies (Chilton et al 1990), it is possible that late-hatched *Z. l. nuttalli* may be exposed to *Z. l. pugetensis* that have already arrived in their winter habitats. It is hard to answer these questions with only the recordings and data of their locality and habitats. However, the strong positive relationship between ecological differences and song in the non-breeding season corresponds with our original hypothesis. Fifth and finally, we did not investigate the other properties of the environment, such as the habitat type, instead focusing only on abiotic climate. Further research on whether the abiotic environment affects the evolution of bird songs directly or indirectly via biotic environmental factors is needed.

In conclusion, using machine learning for automatic analyses, we evaluated divergence within and between two subspecies of the White-crowned Sparrow. Our results indicated that *Zonotrichia leucophrys pugetensis* and *Z. l. nuttalli* have significantly different songs from each other, and both sing different songs in breeding and non-breeding contexts. The divergence in songs correlate with spatial factors such as geographic distance and climate, but subspecies identity and season mediate the response.

## ACKNOWLEDGEMENTS

We thank the Carstens Lab past and present, as well as D. Nicholson and A. Delgado for helpful feedback. We thank two anonymous reviewers, one associate editor, and T. S. Sillett for reviews.

## Funding statement

KLP was funded by NSF DEB-2016189. BCC was funded by NSF DBI-1661029, DEB-2016189, NSF DEB-1831319, and DBI-1910623. JY was funded by the Undergraduate Diversity at Evolution program.

## Author contributions

Experimental design: JY, KLP, BCC. Data collection: JY. Methods development: KLP, BCC. Analyses: JY, KLP. Writing: JY. Figures: JY, KLP. Editing: JY, KLP, BCC.

## Data depository

Data will be available upon acceptance on Dryad. Scripts will be available on GitHub.

## Notes

### Competing Interest Statement

The authors have declared no competing interest.

### Summary of Updates

This version of the manuscript has been revised after peer review. We have added more data, changed our statistical analyses, and updated the text.

