## Supplemental Material for "Machine learning reveals relationships between song, climate, and migration in coastal *Zonotrichia leucophrys*"

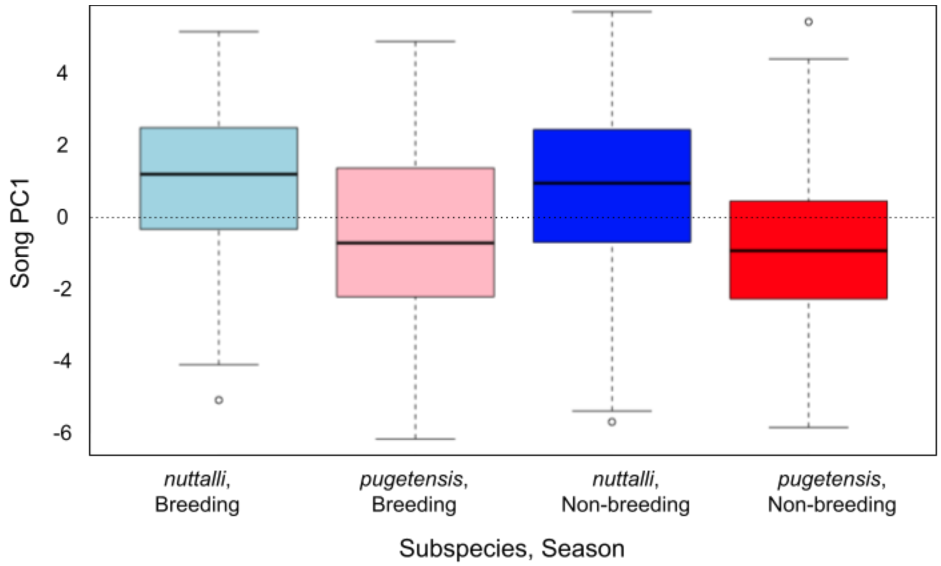


**Supplementary Material Figure S1**: Principal components analysis shows differences between seasons and subspecies for song (PC1).

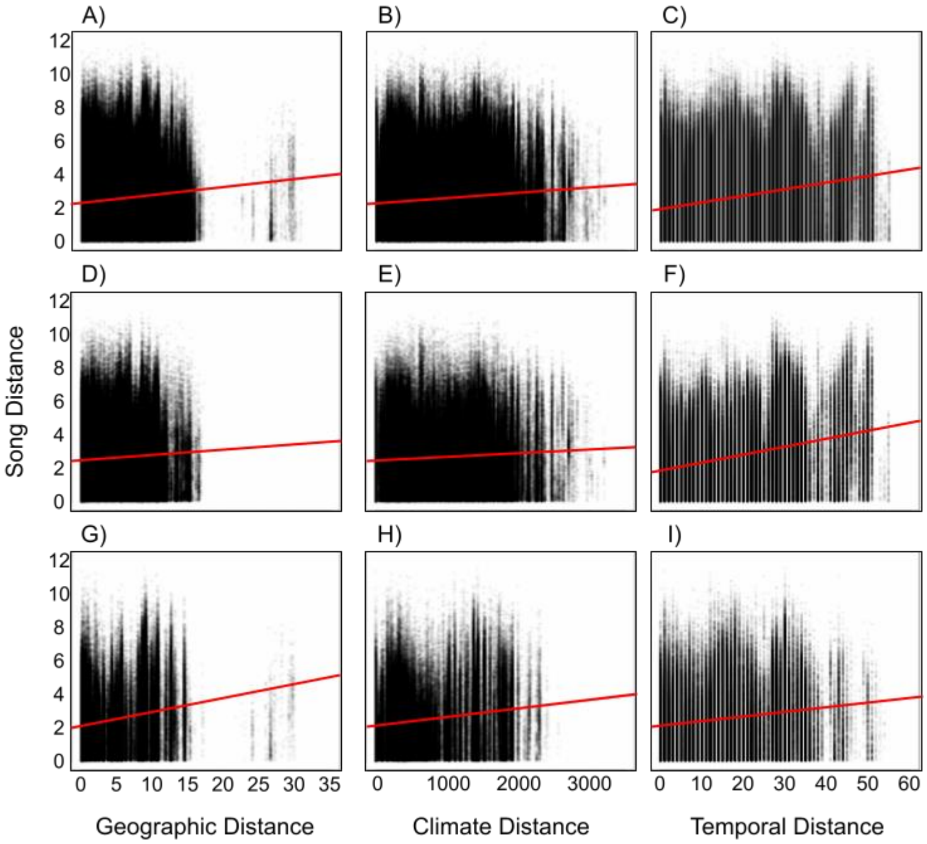


**Supplementary Material Figure S2**. Song distance (y-axis) is correlated with environmental variables all year (A, B, C), in the breeding season (D, E, F), and in the non-breeding season (G, H, I). X-axis depicts geographic distance (A, D, G), climatic distance (B, E, H), and temporal distance (C, F, I). Solid red lines show linear relationship, but all are significant using ANOVAs. Note that individual points are partially transparent to show density of data.

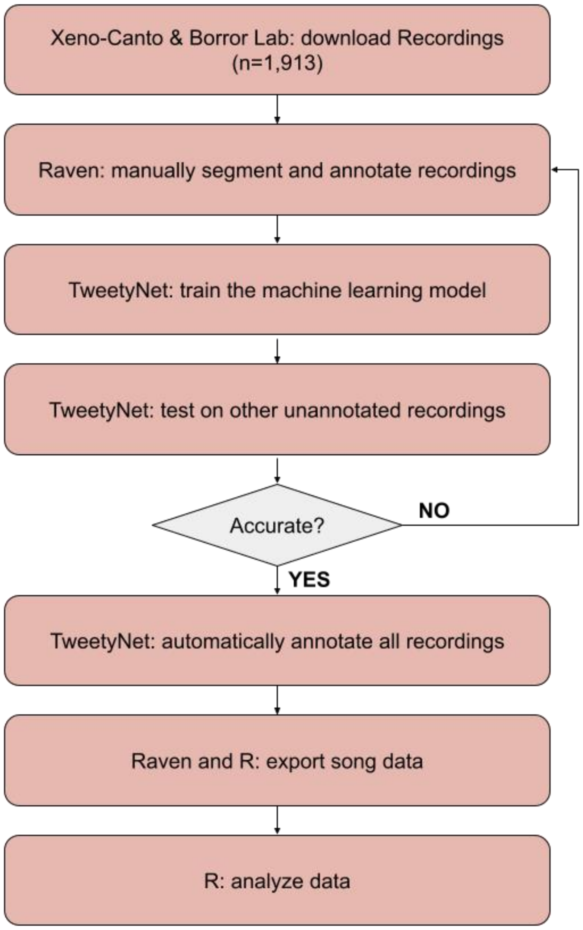


**Supplementary Material Figure S3**. Flowchart showing the process for analysis of data.

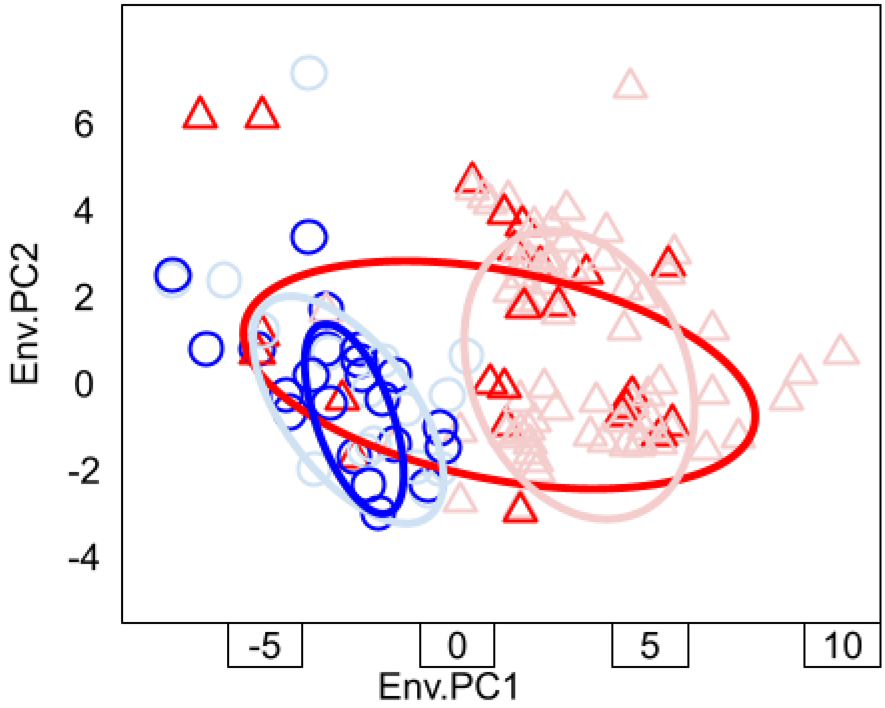


**Supplementary Material Figure S4**. Principal components of the environment for different subspecies in different seasons. X-axis shows Env.PC1, the y-axis shows Env.PC2. The ellipses show 75% confidence. Red triangles represent *pugetensis*. Blue circles represent *nuttalli*. Non-breeding season is in darker shades; breeding season is in lighter shades. Note that some points at the same locality are jittered for visibility.

**Supplementary Material Table S1**: Importance of principal components performed on environmental data. Broken stick refers to expected proportion of variance explained under the broken stick model.

| PC | Standard deviation | Proportion of Variance | Cumulative Proportion | Broken Stick |
|---|---|---|---|---|
| Env.PC1 | 3.467 | 0.633 | 0.633 | 0.187 |
| Env.PC2 | 2.001 | 0.211 | 0.843 | 0.134 |
| Env.PC3 | 1.359 | 0.097 | 0.941 | 0.108 |
| Env.PC4 | 0.846 | 0.038 | 0.978 | 0.090 |
| Env.PC5 | 0.513 | 0.014 | 0.992 | 0.077 |
| Env.PC6 | 0.297 | 0.005 | 0.997 | 0.067 |
| Env.PC7 | 0.172 | 0.002 | 0.998 | 0.058 |
| Env.PC8 | 0.116 | 0.001 | 0.999 | 0.050 |
| Env.PC9 | 0.074 | 0 | 0.999 | 0.044 |
| Env.PC10 | 0.057 | 0 | 1 | 0.038 |
| Env.PC11 | 0.052 | 0 | 1 | 0.033 |
| Env.PC12 | 0.042 | 0 | 1 | 0.028 |
| Env.PC13 | 0.031 | 0 | 1 | 0.023 |
| Env.PC14 | 0.028 | 0 | 1 | 0.019 |
| Env.PC15 | 0.026 | 0 | 1 | 0.016 |
| Env.PC16 | 0.019 | 0 | 1 | 0.012 |
| Env.PC17 | 0.014 | 0 | 1 | 0.009 |
| Env.PC18 | 0.010 | 0 | 1 | 0.006 |
| Env.PC19 | 0 | 0 | 1 | 0.003 |

**Supplementary Material Table S2**: Rotation of Principal Components on Temperature WorldClim Variables only.

| | bio1 | bio2 | bio3 | bio4 | bio5 | bio6 | bio7 | bio8 | bio9 | bio10 | bio11 |
|---|---|---|---|---|---|---|---|---|---|---|---|
| Env.PC1 | -0.273 | -0.200 | -0.192 | 0.114 | -0.209 | -0.223 | -0.038 | -0.250 | -0.227 | -0.235 | -0.251 |
| Env.PC2 | -0.065 | 0.128 | -0.315 | 0.452 | 0.289 | -0.277 | 0.459 | -0.210 | 0.238 | 0.205 | -0.213 |
| Env.PC3 | 0.152 | 0.402 | 0.192 | 0.032 | 0.267 | 0.011 | 0.239 | 0.136 | 0.192 | 0.225 | 0.126 |
| Env.PC4 | 0.244 | -0.402 | -0.295 | 0.052 | 0.018 | 0.327 | -0.210 | 0.190 | 0.324 | 0.316 | 0.177 |
| Env.PC5 | -0.003 | 0.377 | 0.303 | -0.252 | -0.035 | -0.162 | 0.080 | 0.130 | -0.144 | -0.106 | 0.120 |
| Env.PC6 | -0.012 | -0.101 | -0.144 | 0.012 | -0.022 | -0.063 | 0.023 | -0.083 | -0.136 | -0.052 | -0.041 |
| Env.PC7 | 0.066 | 0.036 | 0.202 | 0.087 | 0.409 | 0.589 | -0.032 | -0.406 | -0.146 | -0.171 | -0.194 |
| Env.PC8 | 0.232 | 0.107 | 0.382 | 0.272 | -0.463 | -0.183 | -0.299 | -0.270 | 0.406 | 0.123 | -0.181 |
| Env.PC9 | -0.060 | 0.351 | -0.491 | -0.101 | -0.126 | -0.023 | -0.100 | 0.184 | -0.092 | 0.076 | 0.099 |
| Env.PC10 | -0.280 | -0.290 | 0.320 | 0.251 | 0.126 | 0.014 | 0.106 | 0.479 | 0.014 | -0.063 | -0.170 |
| Env.PC11 | 0.378 | -0.093 | -0.157 | -0.403 | 0.089 | -0.119 | 0.165 | -0.306 | 0.230 | -0.303 | 0.039 |
| Env.PC12 | 0.444 | -0.255 | 0.041 | -0.059 | 0.066 | -0.241 | 0.228 | 0.359 | 0.063 | -0.342 | -0.230 |
| Env.PC13 | 0.178 | 0.402 | -0.245 | 0.244 | -0.079 | 0.242 | -0.240 | 0.293 | 0.044 | -0.335 | -0.400 |
| Env.PC14 | 0.467 | -0.060 | 0.023 | 0.198 | -0.049 | -0.104 | 0.027 | 0 | -0.644 | 0.377 | -0.017 |
| Env.PC15 | 0.070 | 0.096 | -0.101 | 0.014 | 0.024 | 0.079 | -0.033 | 0.043 | -0.024 | -0.086 | -0.109 |
| Env.PC16 | -0.116 | 0.003 | -0.018 | 0.054 | 0.007 | 0.005 | 0.003 | -0.033 | 0.041 | -0.056 | 0.170 |
| Env.PC17 | -0.300 | -0.005 | -0.022 | -0.254 | 0.005 | 0.006 | 0.001 | 0.057 | 0.198 | 0.118 | -0.166 |
| Env.PC18 | 0.019 | 0.007 | -0.018 | 0.477 | -0.020 | -0.029 | 0.001 | -0.016 | 0.025 | -0.444 | 0.660 |
| Env.PC19 | 0 | 0 | 0 | 0 | 0.603 | -0.455 | -0.655 | 0 | 0 | 0 | 0 |

**Supplementary Material Table S3**: Rotation of Principal Components on Precipitation WorldClim Variables only.

| | bio12 | bio13 | bio14 | bio15 | bio16 | bio17 | bio18 | bio19 |
|---|---|---|---|---|---|---|---|---|
| Env.PC1 | 0.250 | 0.231 | 0.262 | -0.261 | 0.238 | 0.271 | 0.270 | 0.229 |
| Env.PC2 | -0.132 | -0.167 | 0.059 | -0.107 | -0.154 | 0.032 | 0.014 | -0.175 |
| Env.PC3 | 0.305 | 0.349 | 0.089 | 0.149 | 0.338 | 0.138 | 0.166 | 0.349 |
| Env.PC4 | 0.023 | -0.055 | 0.328 | -0.143 | -0.034 | 0.263 | 0.247 | -0.062 |
| Env.PC5 | -0.121 | -0.255 | 0.449 | -0.195 | -0.229 | 0.291 | 0.295 | -0.250 |
| Env.PC6 | -0.104 | -0.002 | 0.352 | 0.885 | -0.048 | 0.060 | 0.091 | -0.036 |
| Env.PC7 | -0.114 | 0.127 | 0.310 | -0.113 | -0.127 | -0.068 | -0.126 | 0.071 |
| Env.PC8 | -0.025 | 0.018 | 0.211 | 0.023 | -0.082 | -0.162 | -0.064 | 0.088 |
| Env.PC9 | -0.147 | 0.185 | 0.410 | -0.151 | -0.011 | -0.475 | -0.201 | 0.158 |
| Env.PC10 | 0.028 | -0.082 | 0.159 | -0.009 | -0.021 | -0.492 | 0.317 | 0.077 |
| Env.PC11 | 0.091 | -0.053 | -0.072 | -0.026 | -0.073 | -0.387 | 0.440 | 0.079 |
| Env.PC12 | -0.049 | 0.207 | 0.178 | -0.067 | -0.068 | 0.171 | -0.446 | 0.042 |
| Env.PC13 | 0.044 | 0.071 | -0.255 | 0.081 | -0.100 | 0.102 | 0.287 | -0.150 |
| Env.PC14 | 0.080 | 0.028 | -0.112 | -0.054 | -0.292 | -0.077 | 0.165 | 0.162 |
| Env.PC15 | 0.375 | -0.758 | 0.099 | 0.037 | 0.083 | 0.033 | -0.220 | 0.404 |
| Env.PC16 | 0.758 | 0.175 | 0.098 | 0.042 | -0.412 | -0.102 | -0.170 | -0.351 |
| Env.PC17 | -0.083 | 0.124 | -0.130 | 0.042 | -0.645 | 0.196 | 0.034 | 0.514 |
| Env.PC18 | -0.155 | 0.002 | -0.051 | -0.008 | -0.169 | 0.071 | 0.032 | 0.275 |
| Env.PC19 | 0 | 0 | 0 | 0 | 0 | 0 | 0 | 0 |

**Supplementary Material Table S4**: Differences between season (breeding vs non-breeding) and subspecies (*pugetensis vs nuttalli*) in environmental PCA values occupied using ANOVAs while controlling for both. Headers as in Table 4. Df = 1 for all comparisons. Note that unknown season individuals were removed. SSQ = sum of squares. MSQ = mean sum of squares. F = F-statistic. P = p-value. For these analyses, significant values have p<0.0023. We indicate a near-significant p-value with bold text and a significant value with bold text plus an asterisk. P-values less than 0.001 displayed as 0 for brevity.

| | **Variable** | **Env.PC1** | **Env.PC2** |
|---|---|---|---|
| Mean values | Breeding | 0.167 | -0.146 |
| | Non-breeding | -0.834 | -0.235 |
| | *nuttalli* | -2.753 | -0.631 |
| | *pugetensis* | 2.345 | 0.281 |
| Model parameters: season | SSQ | 467 | 4 |
| | MSQ | 467 | 3.7 |
| | F | 89.11 | 1.327 |
| | P | **0*** | 0.25 |
| Model parameters: subspecies | SSQ | 12399 | 397 |
| | MSQ | 12399 | 397.2 |
| | F | 2363.86 | 142.334 |
| | P | **0*** | **0*** |

**Supplementary Material Table S5**: Differences between season (breeding vs non-breeding) and subspecies (*pugetensis vs nuttalli*) in raw environmental values occupied using ANOVAs while controlling for both. Headers as in Table 4. For these analyses, significant values have p<0.0023. We indicate a near-significant p-value with bold text and a significant value with bold text plus an asterisk. P-values less than 0.001 displayed as 0 for brevity.

| | **Mean values** | | | | **Model parameters: season** | | | | **Model parameters: subspecies** | | | |
|---|---|---|---|---|---|---|---|---|---|---|---|---|
| | **Breeding** | **Nonbreeding** | ***nuttalli*** | ***pugetensis*** | **SSQ** | **MSQ** | **F** | **P** | **SSQ** | **MSQ** | **F** | **P** |
| bio1 | 12.28 | 12.93 | 14.06 | 11.00 | 193 | 193 | 84.34 | **0*** | 4460 | 4460 | 1948.49 | **0*** |
| bio2 | 9.78 | 10.28 | 10.75 | 9.207 | 118 | 118.3 | 55.84 | **0*** | 1136 | 1135.900 | 536.11 | **0*** |
| bio3 | 52.473 | 54.521 | 58.189 | 48.285 | 1950 | 1950 | 79.09 | **0*** | 46765 | 46765 | 1896.59 | **0*** |
| bio4 | 341.93 | 325.84 | 296.16 | 375.77 | 120320 | 120320 | 29.21 | **0*** | 3025158 | 3025158 | 734.39 | **0*** |
| bio5 | 22.682 | 23.227 | 24.032 | 21.745 | 138 | 137.9 | 24.86 | **0*** | 2491 | 2490.900 | 449.02 | **0*** |
| bio6 | 3.926 | 4.311 | 5.506 | 2.614 | 69 | 69 | 41.26 | **0*** | 3991 | 3991 | 2390.99 | **0*** |
| bio7 | 18.756 | 18.916 | 18.525 | 19.132 | 12 | 11.86 | 1.853 | 0.174 | 176 | 175.940 | 27.482 | **0*** |
| bio8 | 8.480 | 9.270 | 10.423 | 7.137 | 290 | 290 | 95.44 | **0*** | 5144 | 5144 | 1692.64 | **0*** |
| bio9 | 16.123 | 16.485 | 17.039 | 15.481 | 61 | 60.9 | 22.44 | **0*** | 1156 | 1156.300 | 426.07 | **0*** |
| bio10 | 16.551 | 16.988 | 17.565 | 15.872 | 89 | 88.8 | 31.24 | **0*** | 1366 | 1366.300 | 480.44 | **0*** |
| bio11 | 8.248 | 9.071 | 10.371 | 6.746 | 315 | 315 | 102.2 | **0*** | 6262 | 6262 | 2034.2 | **0*** |
| bio12 | 1244.6 | 1115.3 | 825.69 | 1569.3 | 7779630 | 7779630 | 33.24 | **0*** | 263666835 | 263666835 | 1126.64 | **0*** |
| bio13 | 219.350 | 201.035 | 170.08 | 254.868 | 155945 | 155945 | 28.47 | **0*** | 3426406 | 3426406 | 625.47 | **0*** |
| bio14 | 15.413 | 11.776 | 2.223 | 26.004 | 6148 | 6148 | 50.7 | **0*** | 269703 | 269703 | 2224.2 | **0*** |
| bio15 | 77.670 | 82.925 | 92.293 | 66.971 | 12843 | 12843 | 144.8 | **0*** | 305691 | 305691 | 3445.7 | **0*** |
| bio16 | 607.68 | 555.928 | 448.257 | 729.045 | 1245304 | 1245304 | 27.42 | **0*** | 37591522 | 37591522 | 827.71 | **0*** |
| bio17 | 63.992 | 50.916 | 11.045 | 107.809 | 79489 | 79489 | 52.79 | **0*** | 4465698 | 4465698 | 2965.77 | **0*** |
| bio18 | 71.729 | 57.719 | 16.782 | 116.825 | 91256 | 91256 | 50.84 | **0*** | 4773348 | 4773348 | 2659.3 | **0*** |
| bio19 | 579.769 | 532.105 | 446.349 | 677.619 | 1056211 | 1056211 | 24.92 | **0*** | 25498358 | 25498358 | 601.55 | **0*** |
